# Non-preferential, but detrimental accumulation of macrophages with clonal hematopoiesis-driver mutations in cardiovascular tissues

**DOI:** 10.1101/2023.09.11.557290

**Authors:** Tsai-Sang Dederichs, Assel Yerdenova, Hauke Horstmann, Tamara Vico, Simone Nübling, Remi Peyronnet, Constantin von zur Muehlen, Timo Heidt, Dennis Wolf, Dirk Westermann, Ingo Hilgendorf

**Affiliations:** Department of Cardiology and Angiology, University Heart Center Freiburg-Bad Krozingen and Faculty of Medicine, University of Freiburg, Freiburg, Germany; Institute for Experimental Cardiovascular Medicine, CardioVascular Biobank, University Heart Center Freiburg-Bad Krozingen and Faculty of Medicine, University of Freiburg, Freiburg, Germany

## Abstract

Clonal hematopoiesis of indeterminate potential (CHIP) is an acquired genetic risk factor for cardiovascular (CV) disease, supposedly mediated by pro-inflammatory recruited monocytes^1–15^. However, how these cells and their progeny behave in the CV tissue remains unclear. Here, we studied human carotid artery plaque and heart tissue samples from *DNMT3A* or *TET2* mutation carriers to quantify the relative accumulation of mutated macrophages and to characterize tissue macrophages from carriers compared to non-carriers. Using droplet digital polymerase chain reaction (ddPCR), we detected similar sizes of CHIP clones in circulating monocytes and macrophages from atheromas and heart tissues, even among CCR2+ (infiltrative), and CCR2– (resident) cardiac macrophages. Using bulk RNA-sequencing (RNA-seq), we revealed a pro-inflammatory gene profile of myeloid cells from CHIP carriers compared to non-carriers. In summary, quantitatively, CHIP mutated myeloid cells did not preferentially accumulate in CV tissues, but qualitatively, they expressed a more disease-prone phenotype.

---

CHIP, defined by somatic mutations in leukemia-associated driver genes, infers a precursor state of hematological malignancies.^1,2^ While CHIP-associated mutations might contribute to genomic instability and skewed clonality^16,17^, they are only mildly leukemogenic. The likelihood for CHIP to develop into overt neoplasia ranges from 0.5% to 1% annually.^9^ In spite of that, while lingering at the precursor state, CHIP significantly precipitates cardiovascular (CV) diseases and death.^8,10–13^ Harboring CHIP is associated with higher risk of coronary artery disease, worse prognosis of congestive heart failure, ischemic stroke, and death.^8,14,15^ Inflammation was revealed in both mouse and human studies as a potential mechanism, by which CHIP transmits CV risk^3,5,8,13^. Studies also showed that circulating monocytes of CHIP carriers are more pro-inflammatory than those of non-carriers,^6,7^ yet to what extend mutated monocyte-derived macrophages accumulate in CV tissues remains unknown. First, we aimed to determine the frequencies of mutated macrophages in human carotid artery plaque and heart tissues relative to those of mutated monocytes in the patients’ blood. Second, we compared the gene expression profiles of atheromatous plaque and cardiac macrophages from CHIP carriers with those from non-carriers.

## Results

### Patient cohorts and immune cells

We screened a total of 28 patients undergoing carotid endarterectomy and 22 patients undergoing heart surgery, among which we identified 9 and 4 CHIP mutation carriers, respectively, **(Extended Data Table 1)** by targeted whole blood DNA sequencing. The average age of the patient group with carotid atherosclerosis was 71.5 ± 11.4 (mean ± standard deviation). More than 80% of them had a carotid stenosis above 70%. The average age of the patient group undergoing heart surgery was 67.1 ± 10.6 (mean ± standard deviation). Nearly 70% of them underwent surgery for heart valve repair with or without additional procedures, five patients were subjected to coronary artery bypass graft surgery alone, and two received a left ventricular assist device. To examine the mutated clone size in the CV tissue and blood, we focused on the carriers (n=4 from each patient group) with the most prevalent CHIP-driver mutations, *DNMT3A* or *TET2*.

From the patients undergoing carotid endarterectomy, we acquired CD14+ CD64+ macrophages from the plaque and CD4+ T cells, CD8+ T cells, CD19+ B cells, and CD14+ monocytes from the blood. From the patients undergoing heart surgery, we acquired CD14+ HLADR– monocytes, and CD14+ HLADR+ CCR2+ and CCR2– macrophages from right atrium appendix or left ventricle as well as CD3+ cells, neutrophils, and CD14+ monocytes from the blood **(Extended Data Figure 1-2)**. DNA and RNA were isolated from the sorted cells for clone size measurement with droplet digital polymerase chain reaction (ddPCR) and transcriptomic profiling with bulk RNA-seq, respectively.

### Comparison of CHIP clone size in blood and CV tissue

As the amount of DNA extracted from sorted macrophages per CV tissue was not sufficient for performing targeted genomic sequencing, we embarked on deploying ddPCR technology, which measures an absolute variant allele frequency (VAF) of predefined targets. For each CHIP carrier, we designed specific primers and probes to target their mutated locus. The primers and probes were first validated using the patients’ whole blood DNA. Owing to the low cell counts we could acquire from the CV tissues, a pre-amplification of the DNA was required to achieve a sufficient number of copies for ddPCR. We therefore first validated the adequacy of the pre-amplification approach using whole blood samples and assured a comparable VAF obtained from targeted genomic sequencing versus ddPCR with or without DNA pre-amplification **(Extended Data Figure 3)**.

The results of ddPCR showed that the frequency of CHIP-mutated monocytes in the blood strongly correlated with that of CHIP-mutated macrophages in both the plaque and the heart with a Pearson correlation coefficient (r) of 0.96 and 0.89, respectively. Both regression lines had a slope close to one, indicating similar frequencies of CHIP mutations in the blood and tissue myeloid cells. The CHIP frequency of monocytes and T cells were weakly correlated with the r values ranging between 0.24 to 0.33. Frequency of CHIP in monocytes and B cells correlated significantly, albeit the slope of the regression line was also as low as 0.21. All in all, we demonstrated that *DNMT3A*- or *TET2*-mutated and non-mutated monocytes had similar capabilities to give rise to macrophages in CV tissues. Low frequency of CHIP clones in blood lymphocytes reflected myeloid skewing driven by *DNMT3A* or *TET2* mutations **(Figure 1A-B)**. In one patient (Carrier 8), we did not detect mutated alleles in the 148 CCR2+ cardiac macrophages. We presume that the absence of mutated cells resulted from low cell counts rather than inability of the CHIP-mutated monocytes to develop into CCR2+ macrophages in the heart, given that we identified a sizable CHIP-mutated clone among CCR2– macrophages (575 cells) in the same sample.

**Figure 1.**
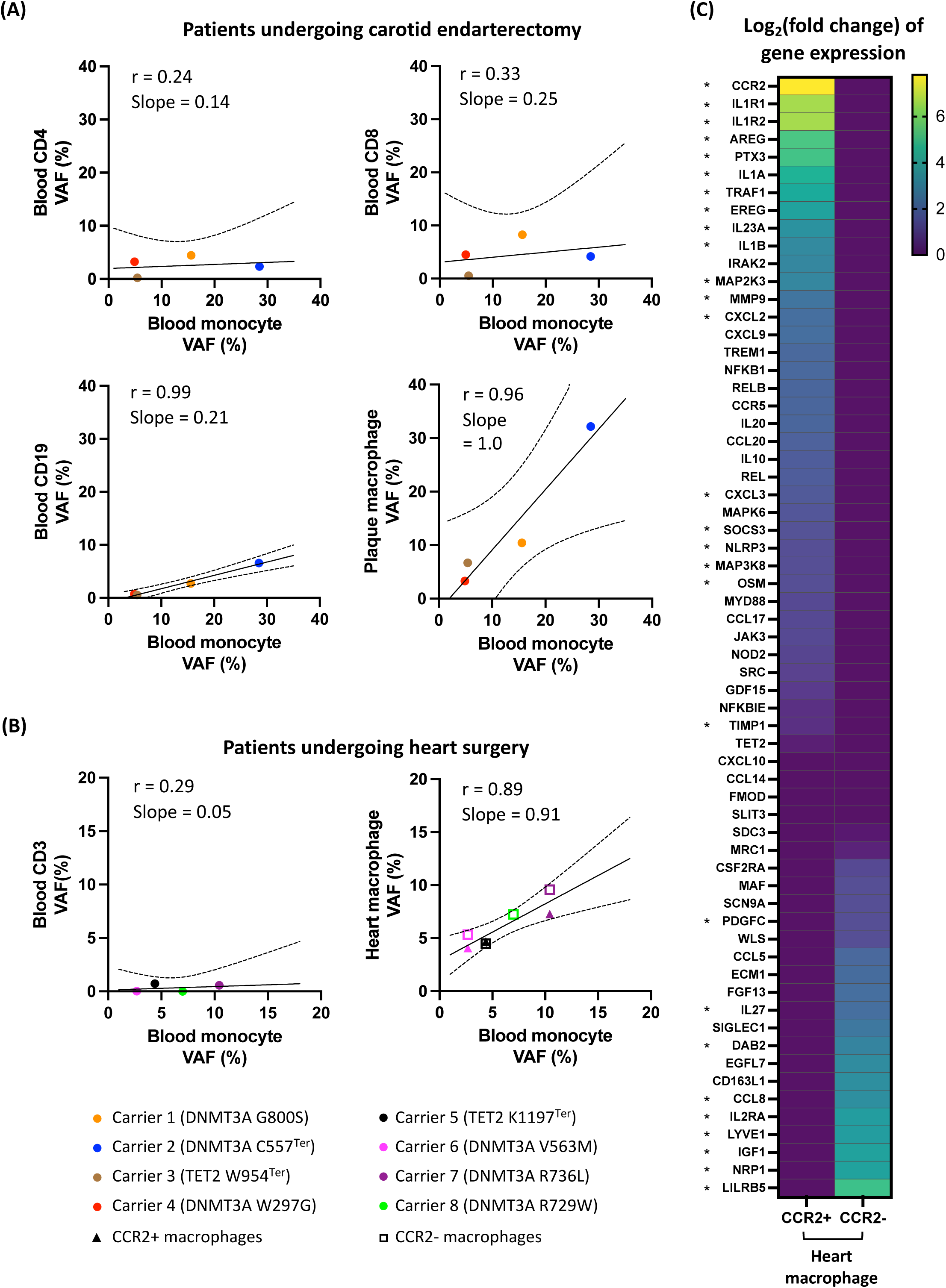
CHIP clone size of immunocytes in blood and CV tissues. **(A-B)** Dot plots depicting CHIP clone size of various cell types versus that of blood monocytes from eight *DNMT3A-* or *TET2-* mutation carriers. Simple linear regression was performed to compute a fit line (solid line) and the 95% confidence bands (dotted lines) for each plot. The Pearson’s correlation coefficient (r) was computed to evaluate correlation. **(B)** A heatmap showing the mean of logarithmic fold change of intra-individual gene expression ratio between the CCR2+ and CCR2– cardiac macrophages. “Ter” indicates translation termination codon, i.e. stop codon. * denotes a *p*-value < 0.05 of two-tailed student’s *t*-test.

Surprisingly, we detected similar sizes of CHIP clones in CCR2+ and CCR2– cardiac macrophages **(Figure 1B right)**, challenging the current paradigm that CCR2– cardiac macrophages represent a resident population with minimal monocyte contribution.^18^ To inspect the distinct gene expression profiles of CCR2+ and CCR2– macrophages in the human heart, we explored the gene signatures published by Bajpai and colleagues^18^ and confirmed the upregulation of pro-inflammatory genes in our sorted CCR2+ macrophages (eg. *NLRP3* and *IL1B*) as well as the upregulation of negative immunomodulators and so-called tissue resident macrophage markers in our sorted CCR2– macrophages (eg. *LILRB5* and *LYVE1*) **(Figure 1C)**. Our finding thus suggests that in the elderly, CCR2+ and CCR2– cardiac macrophages have been repopulated by blood monocytes at comparable rates.

### Gene expression of myeloid cells from CHIP mutation carriers

Although *DNMT3A*- or *TET2*-mutated macrophages did not dominate accumulation in the CV tissue, we suppose that dysregulated immune activities of these cells still play a role in enforcing the associated CV risks. To evaluate whether tissue macrophages from the *DNMT3A*- or *TET2*- mutation carriers present distinctive gene profiles, we selected 16 non-carriers (eight from each patient group) as controls, matched for sex, age, body mass index, disease severity, blood work, and medical histories **(Extended Data Table 2-3)**. Bulk RNA-seq revealed an upregulation of genes associated with active immune responses in myeloid cells from the *DNMT3A-* or *TET2-* mutation carriers, especially in tissue macrophages **(Figure 2A-C)**. Many of these differentially expressed genes (DEG) promote inflammation and chemotaxis. Interestingly, although CCR2+ macrophages generally exhibited a more pro-inflammatory profile compared to their CCR2– counterparts **(Figure 1C)**, our data revealed that both macrophage subsets from CHIP carriers displayed an overall higher expression of inflammatory genes (e.g., IL1B, BIRC3, and CCL3), as opposed to the corresponding macrophages from the non-carriers **(Figure 2C, D)**. These results suggest a widespread activation of myeloid cells resulting from *DNMT3A-* and *TET2-*mutations.

**Figure 2.**
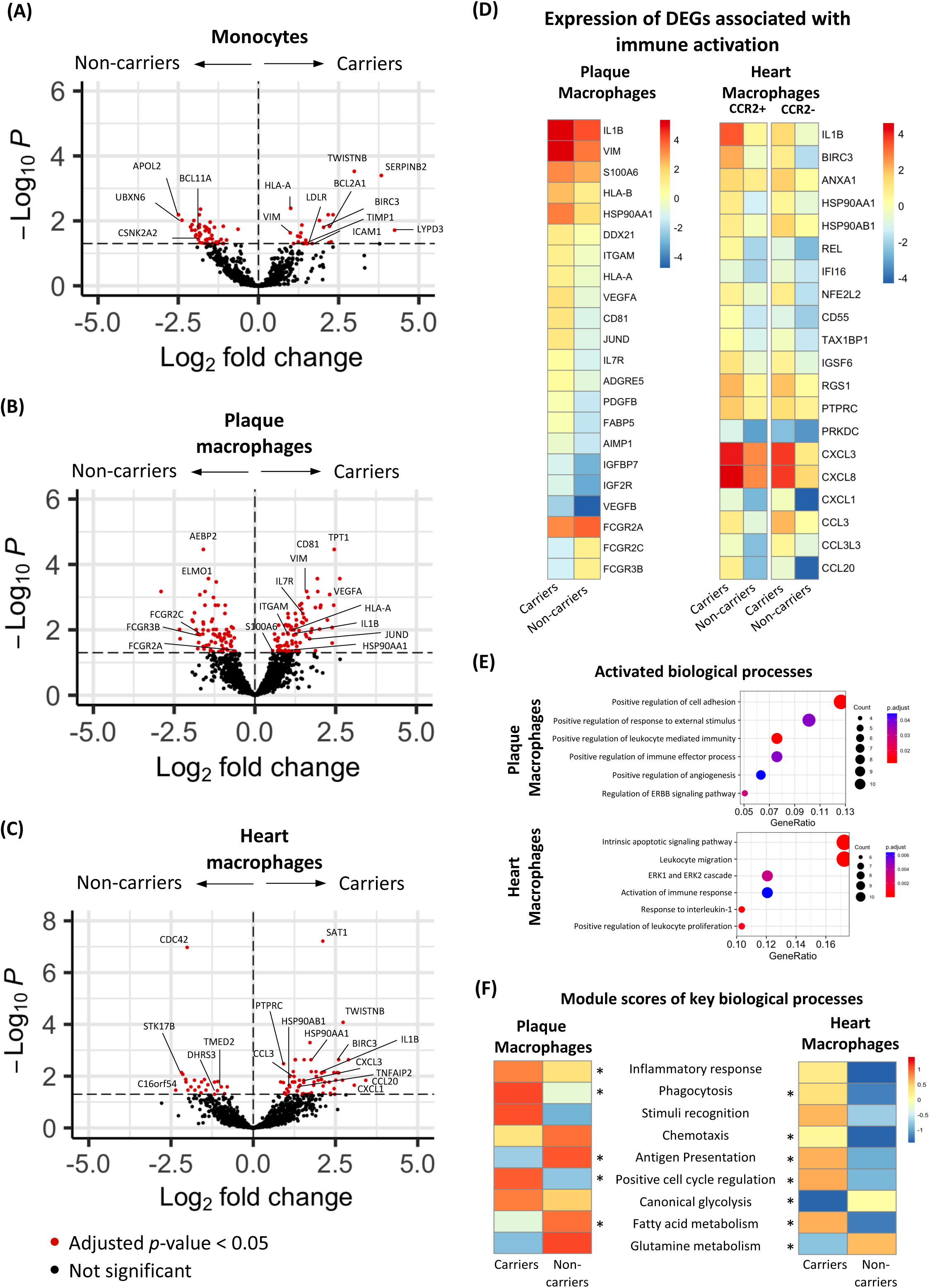
Activated immune responses of tissue macrophages from *DNMT3A-* or *TET2-*mutation carriers. **(A-C)** Volcano plots showing DEGs (red dots) between carriers and non-carriers. **(D)** Heatmaps of representative DEGs associated with activated immune responses. **(E)** Dot plots of enriched gene ontology (GO) terms of upregulated DEGs in plaque and heart macrophages from CHIP mutation carriers. **(F)** Heatmaps depicting the mean of module scores of plaque and heart macrophages from carriers and non-carriers. * denotes *p*-values < 0.05 of one-tailed Mann-Whitney test.

While in general inflammation was a prominent phenotype of both plaque and cardiac macrophages in *DNMT3A-* and *TET2-*mutation carriers, some enriched biological processes by the upregulated DEGs differed in the plaque and heart macrophages **(Figure 2E)**. Module scores, which quantitatively summarized specific biological processes, demonstrated that macrophages from CHIP carriers showed stronger inflammatory responses and phagocytosis activity, along with a positive regulation of cell cycle in both plaque and heart tissue. However, CHIP mutations were associated with an enhanced chemotaxis activity and antigen presentation of macrophages isolated from the heart, while gene expression associated with these immune functions were relatively reduced in plaque macrophages from the CHIP carriers. CHIP mutations also impacted genes linked to macrophage metabolism suggestive of enhanced energy production by fatty acid consumption in cardiac macrophages. On the contrary, plaque macrophage gene signatures of CHIP carriers pointed towards more glycolysis and less fatty acid or glutamine metabolism. **(Figure 2F, Extended Data Figure 4)**

## Discussion

Our work showed a CHIP prevalence of 32 % in patients with overt carotid atherosclerosis and 18% in patients undergoing heart surgeries. Despite variations in CHIP screening techniques and patient age across different studies, these rates are similar to those observed in patients with chronic heart failure^14^, aortic valve stenosis^19^, and those at risk of coronary artery diseases^20^.

Accumulation of pro-inflammatory macrophages in the CV tissue is one of the key drivers of atherosclerosis and heart disease.^21^ Plaque macrophages, in particular, derive from infiltrating monocytes that proliferate locally as disease advances^22,23^. Thus, we hypothesized that CHIP-associated CV risk was linked to augmented infiltration or local proliferation of CHIP-mutated monocytes. While our research marks the initial evidence of CHIP clones’ presence in CV tissue, the comparable CHIP clone size between blood monocytes and tissue macrophages suggested an equivalent net accumulation of mutated and non-mutated monocytes and their progeny in the CV tissue. The significantly smaller CHIP clones in the lymphocyte population reinforce the phenomenon of CHIP-driven myeloid skewing. As a noticeable frequency of mutated B cells were formerly reported in some patients carrying *TET2* and *DNMT3A* mutations^17,24^, we identified a steady 1:5 ratio of mutated B cells and monocytes in the blood.

Another distinctive aspect of our research lies in the identification of significant CHIP clones within CCR2– cardiac macrophages. These macrophages are currently labeled as tissue-resident macrophages of prenatal origin. Circulating monocytes were reported to contribute trivially to the CCR2– cardiac macrophage population in patients receiving sex-mismatched heart transplantation^18^. However, given that CHIP mutations are acquired only at a relatively advanced age, i.e. older than 40 years, we suppose that these CHIP-mutated CCR2– cardiac macrophages were derived from infiltrating monocytes originated from the bone marrow. We also confirmed that these CCR2– macrophages expressed a reparative, anti-inflammatory phenotype as shown previously^18^. Our data thus support using CCR2 to identify reparative tissue macrophages but cast doubt on using CCR2 as a single marker to distinguish the origins of tissue macrophages in humans. Our data align with a recent report of CHIP mutations in brain microglia which are also believed to maintain independent from monocytes.^25^

Although we did not observe an enrichment of CHIP-mutated myeloid cells in the CV tissue, we discovered that the circulating and tissue myeloid cells from *DNMT3A* or *TET2* mutation carriers present a distinctive transcriptome as opposed to those from the matched non-carriers. In the plaque, macrophages from the carriers upregulated genes promoting inflammation, immune responses, cellular differentiation and apoptosis. Similar immune activation phenotypes were shown in the heart macrophages with additional chemokines **(Figure 2A-D)** upregulated in the carriers. Many of these upregulated genes can function intracellularly to activate macrophages or be released to the extracellular space to amplify inflammatory activities through other cells^26–28^. On the other hand, our data also showed distinctive impacts of CHIP on plaque and cardiac macrophage functions and metabolism. These distinctions could result from the tissue type where macrophages adapt to their local environment or the general disease context. The formation of carotid artery plaques involves excessive lipid deposition, endothelial damage, and inflammatory immunocyte infiltration at the lesions^29–31^, while heart diseases encompass malfunctioning cardiomyocytes, fibroblasts and imbalanced immune activities in response to intrinsic or extrinsic (e.g. ischemia, valvular defects) triggers.^32–34^ Overall, our data manifest that CHIP mutations associate with inflammatory macrophages in both CV tissues.

A limitation of our study is that the bulk-RNA sequencing technique we employed, does not discriminate between mutated and non-mutated cells. Therefore, we cannot comment on whether the inflammatory signature of tissue macrophages in CHIP carriers solely derived from the relatively small fraction of mutated clones or whether the majority of non-mutated macrophages contributed as well, for example following paracrine activation. Addressing this open question will require single-cell multiomic techniques to uncover a direct connection between the cellular genotypes and phenotypes per cell.

## Conclusion

Our study manifested the presence of pro-inflammatory myeloid cells in human plaque and heart tissues from CHIP mutation carriers. The equivalent size of CHIP clones in the circulation and tissue indicate a comparable ability of mutated and non-mutated monocytes to accumulate and differentiate into macrophages in CV lesions. Similar mutation frequencies among circulating monocytes and both CCR2+ and CCR2– cardiac macrophages suggest that, contrary to the current paradigm, CCR2– macrophages in the aged human heart derive from infiltrating monocytes. While CHIP-mutated macrophages did not enrich locally, their gene expression profile reinforced that inflammation contributes to CHIP-associated CV disease and may represent a therapeutic target.

## Methods

### Study cohorts and workflow

We conducted an observational study in patients undergoing carotid endarterectomy and heart surgery at the University Hospital Freiburg. The study complies with the Declaration of Helsinki and was approved by the local ethics committee (EK 249/14, 393/16, 6/20). With written informed consent, patients’ blood and CV tissue retrieved during the surgeries were immediately processed for fluorescence-activated cell sorting (FACS) of lymphocytes and myeloid cells. **(Extended Data Figure 1)**. An aliquot of whole blood was subjected to targeted genomic sequencing for CHIP mutation screening.

### CHIP mutation screening

Genomic DNA (gDNA) of 350 μL citrated whole blood was extracted using DNeasy Blood and Tissue Kit (QIAGEN). Fifty nanogram of total gDNA of each sample was constructed into a targeted genomic library according to the instruction of the TruSight Myeloid Sequencing Panel (Illumina) and sequenced either on a MiSeq instrument with 220 bp paired-end reads or on a NextSeq500 instrument with 150 bp paired-end reads. Each sample was sequenced with a mean depth of 1M to 4M reads. Data analysis was performed on the Galaxy Europe platform (usegalaxy.eu). In brief, we first applied Trim Galore! (v0.4.3.1) to remove adapters, low-quality reads (Phred score < 28), and 22 bp from the 5’ end of both read 1 and read 2. The trimmed reads were mapped to the human genome assembly GRCh37 (hg19) using bowtie2 (v2.3.4.3). After converting bam files to pileup using Generate pileup (v.1.1.3), we identified variants at positions having a base quality of Phred score > 28 with VarScan (v2.4.2) according to the consensus genotype. The variants were subsequently annotated with SnpEff eff (v4.3) and GEMINI (v0.20.1). Confirmed CHIP variants should be covered with a read depth (DP) > 2500 and carry a variant allele frequency (FREQ) above 2% on exonic regions. We ruled out variants with a FREQ of 40% to 55% or above 90%, or those occur in more than 0.5% of non-Finnish European population to exclude germline mutations. Variants fulfilling the following criteria were considered technical artifacts and thus excluded: 1. Having a discrepancy of FREQ above 50% between forward and reverse reads. 2. Occurring in more than 10% of the samples of the same library. 3. Locating on the oligo positions of the TruSight Myeloid Sequencing Panel. 4. Occurring only in a specific amplicon at the position where multiple amplicons overlap.

### Blood and tissue sample preparation

Ten mL citrated whole blood from each patient first underwent red blood cell lysis for 10 minutes and washed with FACS buffer (Ca/Mg2+ free phosphate buffered saline (PBS) + 5% fetal calf serum). Aortic plaque or heart samples were shredded into fine pieces by mechanical forces and incubated in digestion mixes at 37 °C **(Details listed in Extended Data Table 4)**. The digested tissue were filtered through strainers and washed with FACS buffer to retrieve single cell suspensions. Thereafter, the cells were stained with the corresponding FACS panels **(Extended Data Table 5)** at 4 °C for 30 minutes and sorted using BD Aria III (BD Biosciences) or MoFlo Astrios EQ (Beckman Coulter) following the gating strategy presented in **Extended Data Figure 2**. The cells were directly deposited in DNA LoBind tubes (Eppendorf) containing buffer RLT Plus (QIAGEN) with 1% β-Mercaptoethanol (PanReac AppliChem) and stored at - 20 °C. DNA and RNA of sorted cells were extracted using AllPrep DNA/RNA micro kit (QIAGEN) according to the manufacturer’s instruction. DNA amount of blood cells was measured using high-sensitivity Qubit dsDNA assay (ThermoFisher), while DNA amount of tissue cells were extrapolated based on the sorted cell counts.

### ddPCR

Isolated DNA of sorted cells from the *DNMT3A* or *TET2* mutation carriers were subjected to ddPCR to measure the CHIP clone size in each sorted cell type. Individual primers were designed with Primer3 (v4.1.0) and first validated with whole blood samples to confirm the annealing temperature. Probes were designed with BeaconDesigner (v7.0) **(Extended Data Figure 3A)** and ordered from Integrated DNA Technologies. Genomic DNA isolated from sorted cells were pre-amplified to the target value of 6 to 24 billion number of copies and subsequently purified using Agencourt AMPure XP beads (Beckman Coulter). The purified DNA were compartmented into oil droplets together with ddPCR Supermix (Bio-rad) as well as customized primers and probes using QX200 droplet Generator (Bio-rad) and underwent another 40 cycles of PCR reaction to enhance the fluorophore signal in the droplets. The samples were read and analyzed using QX200 ddPCR Droplet Reader and QuantaSoft software (Bio-Rad). We purchased all ddPCR consumables from Bio-Rad and followed the instructions of the QX200 ddPCR system.

### Bulk RNA-seq and data analysis

RNA of the carriers and non-carriers were constructed into transcriptome libraries using NEBNext Single Cell/Low Input RNA Library Prep Kit (New England BioLabs) and sequenced using NextSeq2000 (Illumina) instrument with 50 bp paired-end reads. The sequenced raw data were first processed into count matrices on the Galaxy Europe platform. In short, we first applied Trim Galore! (v0.4.3.1) to remove adapters and low-quality reads (Phred score < 28) and mapped the trimmed reads to human genome assembly GRCh38 (hg38) using RNA STAR (v2.7.10). The gene expression was quantified using featureCounts (v2.0.3). We then further analyzed the gene expression matrices in R (v4.1). After removal of ribosome genes, the rest of genes with total counts between 5,000 to 500,000 among all samples were used to compute DEGs using DESeq2 (v1.40.1). The biological process enrichment was analyzed using clusterProfiler (v4.8.1). The Volcano plots were generated using EnhancedVolcano (v1.18.0). Module scores were calculated using all genes categorized in the specific gene ontology (GO) terms **(Extended Data Table 6)**.

## Acknowledgement

We thank Kristina Kollmar, the technical assistant of CVBB of Institute for Experimental Cardiovascular Medicine, and Markus Lengemann for their support in patient heart tissue collection. We thank the staff and patients of the Department for Cardiovascular Surgery of the University Heart Centre Freiburg - Bad Krozingen for access to human tissue. We thank staff members of the Lighthouse core facility at University of Freiburg, Marie Follo, Jan Bodinek-Wersing, and Dieter G. Herchenbach, for their help and guidance with cell sorting and ddPCR. We thank Dietmar Pfeifer, head of genomics laboratory of University Hospital Freiburg, for his support in CHIP mutation screening and establishing ddPCR methodology. We also thank Vladimir Benes, Head of Genomic Core EMBL, Heidelberg, Björn Grüning, head of Freiburg Galaxy Team, and Rolf Backofen, head of Bioinformatics Group Freiburg, and the SFB1425 S3 service project for their help and advice with sequencing. We thank Klaus Kaier for his support in statistical analysis.

## Author Contributions

TD, AY, and HH collected and processed patient samples. TD and AY analyzed clinical data, established and performed experiments. RP and SN coordinated cardiovascular biobank (CVBB), supported patient sample and clinical data collection. TD, DWo, and IH designed the study. IH acquired funding. TD and IH analyzed and interpreted the data. The original manuscript was drafted by TD and IH, and reviewed/edited by TV, TH, CM, DWo, and DWe.

## Data availability

The RNA-seq data are deposited in the Gene Expression Omnibus (GEO) DataSets with the accession number of GSE226642.

## Extended Data

**Extended Data Figure 1.**
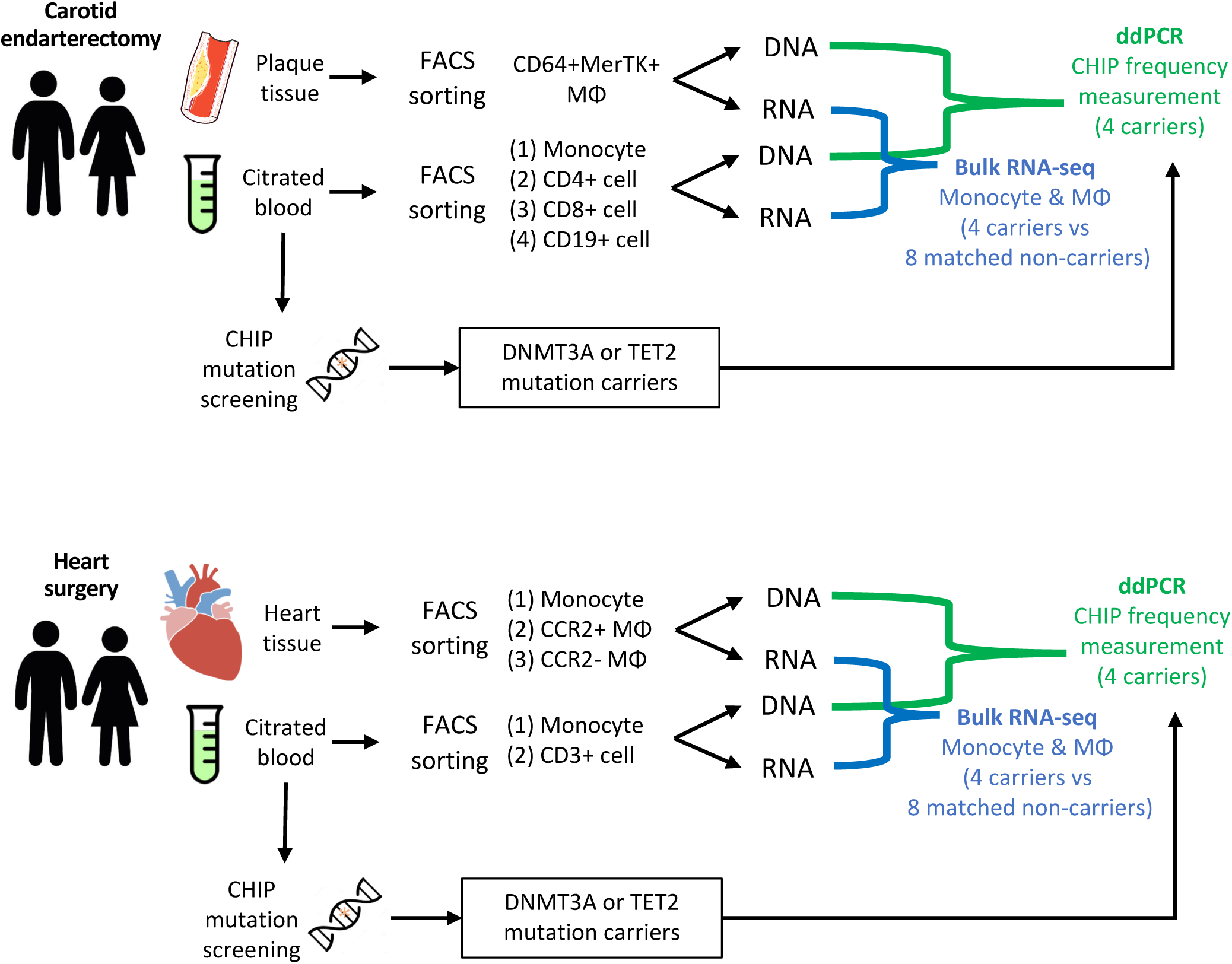
Study design and workflow. Schematic illustration of the study workflow.

**Extended Data Figure 2.**
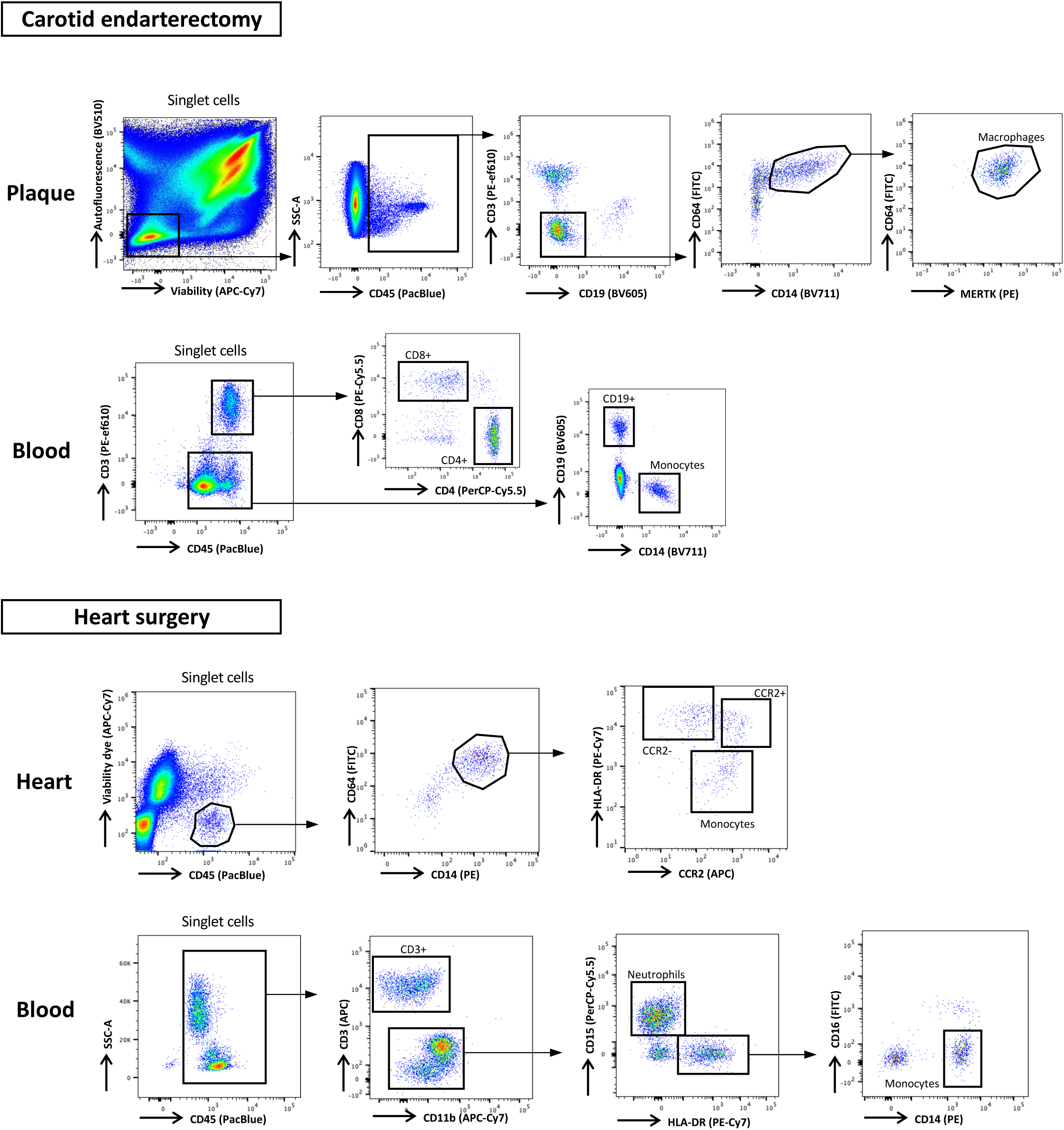
FACS gating strategy of blood and tissue samples. FACS dot plots illustrating the gating strategy of myeloid cells and lymphocytes from blood and tissue samples in the two patient cohorts.

**Extended Data Figure 3.**
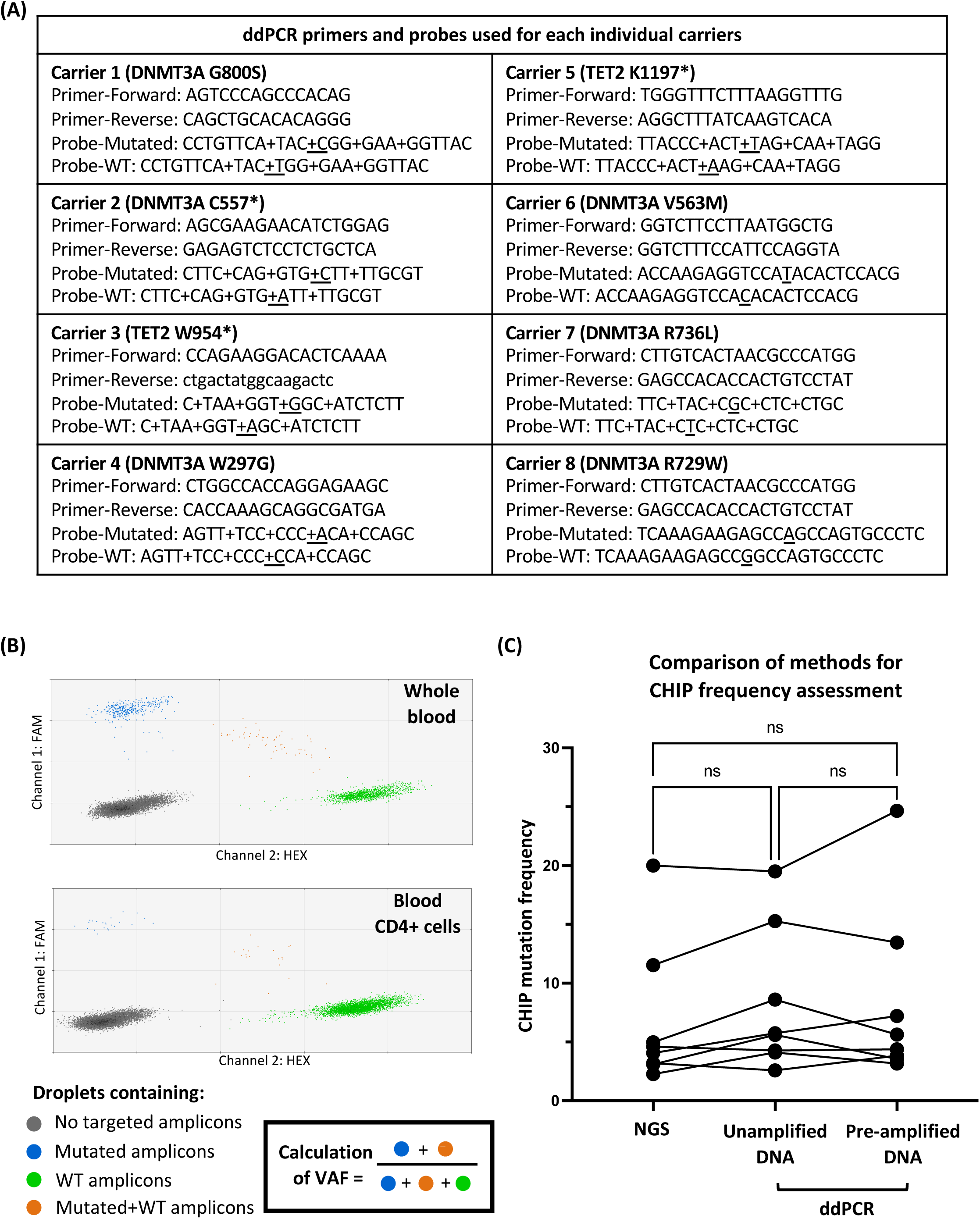
ddPCR method establishment and validation. **(A)** Oligonucleotide sequence of primers and probes targeting specific mutated site of each carrier. Probes targeting mutated and wild-type (WT) gDNA were labeled with FAM and HEX, respectively. + denotes locked nucleic acids. Underlined nucleotide in probes indicates the single nucleotide polymorphism of the carriers. **(B)** Schematic illustration of how variant allele frequency (VAF) was calculated based on the ddPCR results. Adequate ddPCR results with at least one double-negative cluster and one single-positive cluster in the two-dimensional figure were selected for gating and droplet counting. Each sample had at least three technical replicates. **(C)** A dot plot comparing the CHIP mutation frequency in the whole blood measured by three methods, namely next generation sequencing (NGS), ddPCR using unamplified DNA, and pre-amplified DNA. The lines connect dots of the same carriers. ns denotes no significant differences between the three methods evaluated by repeated measures one-way ANOVA with the Geisser-Greenhouse correction and Turkey’s multiple comparisons test.

**Extended Data Figure 4.**
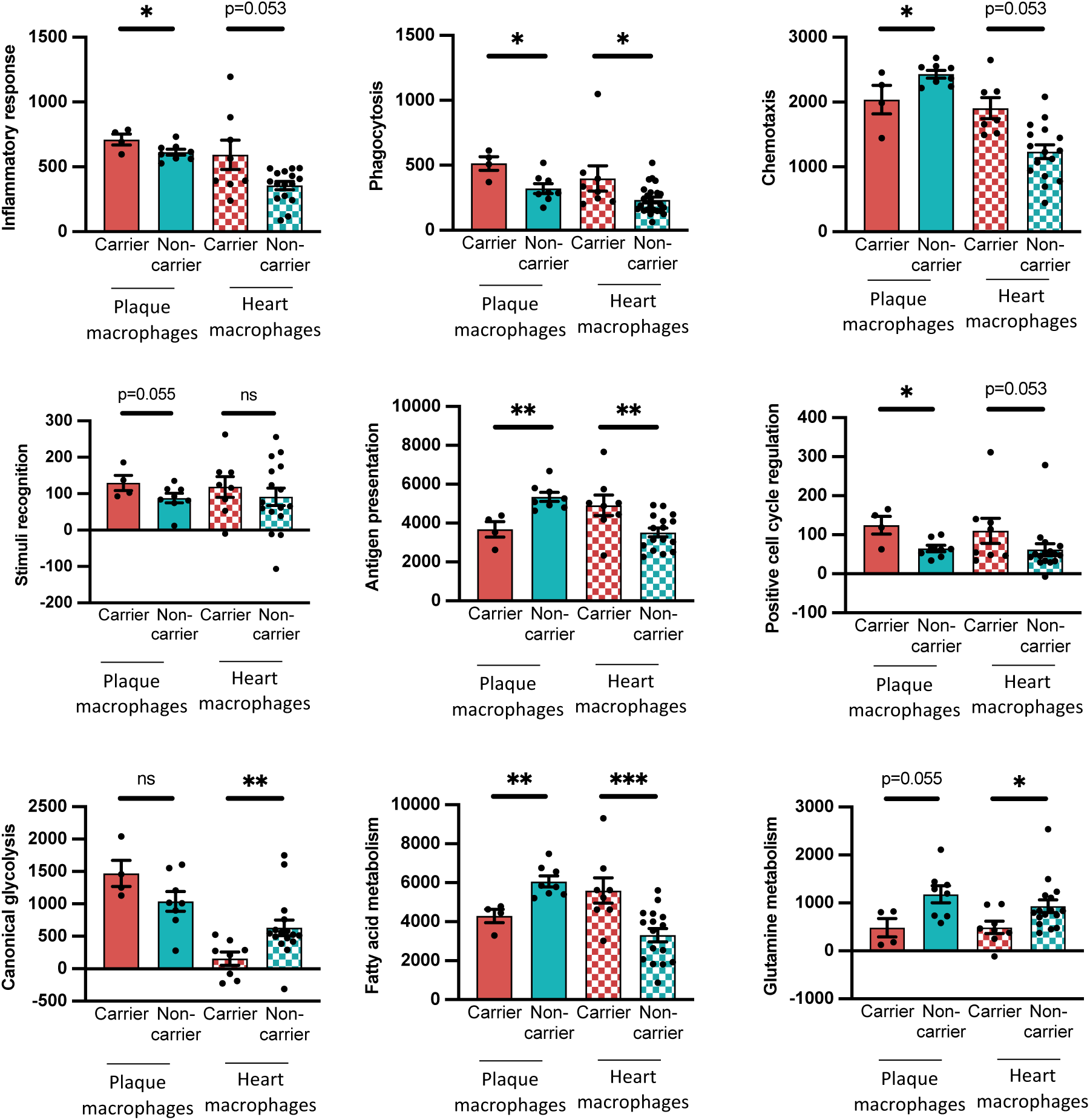
Module scores of biological processes associated with immune functions. Bar charts showing module scores of selected biological processes of tissue macrophages from carriers and non-carriers. Genes comprising the specific GO pathways **(Extended Data Table 6)** were used for module score calculation. The *p*-values were calculated using one-tailed Mann-Whitney test. ns denotes no significant differences, * denotes *p*-value ≤ 0.05, ** denotes *p*-value ≤ 0.01, *** denotes *p*-value ≤ 0.001.

**Extended Data Table 1.** CHIP mutations of patients undergoing carotid endarterectomy or heart surgery CHIP mutations identified among patients who underwent carotid endarterectomy.

| POS | FREQ | chrom | ref | alt | rs_ids | gene | exon | aa_change | impact |
| --- | --- | --- | --- | --- | --- | --- | --- | --- | --- |
| 209113112 | 19.42% | chr2 | C | T | rs121913500 | IDH1 | 4 | R132H | missense_variant |
| 36932358 | 2.35% | chr1 | T | C | None | CSF3R | 17 | K731R | missense_variant |
| 25462009 | 11.54% | chr2 | C | T | rs757083492 | DNMT3A | 20 | G800S | missense_variant |
| 25467204 | 20% | chr2 | G | T | None | DNMT3A | 15 | C557* | stop_gained |
| 209113112 | 18.88% | chr2 | C | T | rs121913500 | IDH1 | 4 | R132H | missense_variant |
| 33792731 | 37.02% | chr19 | G | GGCGGGT | rs762459325 | CEBPA | 1 | P232HPP | inframe_insertion |
| 106157960 | 2.27% | chr4 | G | A | None | TET2 | 3 | W954* | stop_gained |
| 36231771 | 3.66% | chr21 | T | C | None | RUNX1 | 6 | R205G | missense_variant |
| 25470585 | 4.61% | chr2 | A | C | rs751916447 | DNMT3A | 8 | W297G | missense_variant |

**CHIP mutations identified among patients who underwent heart surgery**
| POS | FREQ | chrom | ref | alt | rs_ids | gene | exon | aa_change | impact |
| --- | --- | --- | --- | --- | --- | --- | --- | --- | --- |
| 106164079 | 3.19% | chr4 | A | T | None | TET2 | 5 | K1197* | stop_gained |
| 25467188 | 3.09% | chr2 | C | T | None | DNMT3A | 15 | V563M | missense_variant |
| 25463286 | 4.06% | chr2 | C | A | None | DNMT3A | 19 | R736L | missense_variant |
| 25463308 | 4.98% | chr2 | G | A | rs200018028 | DNMT3A | 19 | R729W | missense_variant |

POS, position; FREQ, variant allele frequency; chrom, chromosome; ref, reference nucleotide; alt, alternative nucleotide; rs\_ids, Single Nucleotide
Polymorphism identification number; aa\_change, change of amino acid.

**Extended Data Table 2.** Clinical characteristics of patients undergoing carotid endarterectomy.

|  | Carrier (n=4) | Non-carrier (n=8) | *p*-value |
| --- | --- | --- | --- |
| Sex [no.] (male:female) | 3:1 | 5:3 | >0.99 |
| Age [yrs] Mean (SD) | 78.75 (11.90) | 67.13 (12.02) | 0.28 |
| Body mass index [kg/m<sup>2</sup>] Mean (SD) | 26.67 (4.41) | 31.17 (10.32) | 0.83 |
| Side of endarterectomy [no.] (left:right) | 2:2 | 5:3 | >0.99 |
| Degree of carotid artery stenosis [%] Mean (SD) | 75 (7.07) | 76 (6.9) | >0.99 |
| **Blood work** |  |  |  |
| Leukocyte [10<sup>3</sup>/μL] Mean (SD) | 8.65 (4.04) | 8.85 (3.80) | 0.81 |
| Neutrophil [10<sup>3</sup>/μL] Mean (SD) | 6.29 (3.41) | 5.97 (2.62) | 0.78 |
| Lymphocyte [10<sup>3</sup>/μL] Mean (SD) | 1.6 (0.57) | 1.87 (0.79) | 0.54 |
| Monocyte [10<sup>3</sup>/μL] Mean (SD) | 0.66 (0.25) | 0.70 (0.19) | 0.82 |
| Thrombocyte [10<sup>3</sup>/μL] Mean (SD) | 238.3 (58.07) | 278.5 (85.11) | 0.57 |
| Hemoglobin [g/dL] Mean (SD) | 11.53 (2.46) | 12.28 (1.19) | 0.77 |
| Hematocrit [%] Mean (SD) | 33.18 (6.75) | 35.16 (3.71) | 0.81 |
| Hemoglobin A1c [%] Mean (SD) | 5.98 (0.21) | 6.08 (0.58) | 0.69 |
| Cholesterol [mg/dL] Mean (SD) | 115.3 (22.32) | 150.5 (38.49) | 0.05 |
| Homologous LDL [mg/dL] Mean (SD) | 52.50 (20.98) | 85.75 (29.73) | 0.07 |
| Homologous HDL [mg/dL] Mean (SD) | 51.25 (7.4) | 51.75 (14.0) | 0.90 |
| Lipoprotein(a) [mg/dL] Mean (SD) | 42.50 (34.65) | 63.33 (52.06) | 0.85 |
| Triglyceride [mg/dL] Mean (SD) | 105.8 (36.34) | 119.3 (53.95) | 0.71 |
| **Medical history** |  |  |  |
| Coronary artery disease [no.] (%) | 1 (25) | 3 (37.5) | >0.99 |
| Hypertension [no.] (%) | 3 (75) | 6 (75) | >0.99 |
| Diabetes mellitus [no.] (%) | 0 (0) | 2 (25) | 0.52 |
| Transient ischemic attack [no.] (%) | 1 (25) | 1 (12.5) | >0.99 |
| Stroke [no.] (%) | 3 (75) | 4 (50) | 0.58 |
| Use of statins [no.] (%) | 4 (100) | 8 (100) | >0.99 |

Continuous variables are shown as mean (SD); significance tested using two-sided Mann-Whitney test.
Categorical variables are shown as frequency (%); significance tested using Fisher's exact test.
HDL, high-density lipoprotein; LDL, low-density lipoprotein.

**Extended Data Table 3.** Clinical characteristics of patients undergoing heart surgery.

|  | Carrier (n=4) | Non-carrier (n=8) | *p*-value |
| --- | --- | --- | --- |
| Sex [no.] (male: female) | 3:1 | 4:4 | 0.58 |
| Age [yrs] Mean (SD) | 69.50 (5.97) | 61.50 (11.77) | 0.25 |
| Body mass index [kg/m<sup>2</sup>] Mean (SD) | 27.78 (6.20) | 30.07 (4.54) | 0.56 |
| Left ventricular ejection fraction [%] Mean (SD) | 0.50 (0.10) | 0.52 (0.13) | 0.37 |
| Tissue source [no.] (left ventricle: right atrium) | 1:3 | 2:6 | >0.99 |
| **Blood work** |  |  |  |
| Leukocyte [10<sup>3</sup>/μL] Mean (SD) | 6.81 (0.57) | 8.14 (1.95) | 0.21 |
| Thrombocyte [10<sup>3</sup>/μL] Mean (SD) | 204.5 (39.23) | 262.1 (98.19) | 0.23 |
| Hemoglobin [g/dL] Mean (SD) | 13.23 (1.31) | 13.83 (1.48) | 0.43 |
| Hematocrit [%] Mean (SD) | 39.50 (3.67) | 40.83 (4.88) | 0.79 |
| **Medical history** |  |  |  |
| Coronary artery disease [no.] (%) | 1 (25) | 3 (37.5) | >0.99 |
| Hypertension [no.] (%) | 1 (25) | 3 (37.5) | >0.99 |
| Diabetes mellitus [no.] (%) | 2 (50) | 1 (12.5) | 0.23 |
| Transient ischemic attack [no.] (%) | 0 (0) | 0 (0) | >0.99 |
| Stroke [no.] (%) | 0 (0) | 0 (0) | >0.99 |
| Use of statins [no.] (%) | 2 (50) | 4 (50) | >0.99 |

Continuous variables are shown as mean (SD); significance tested using two-sided Mann-Whitney test.
Categorical variables are shown as frequency (%); significance tested using Fisher's exact test.
Comparison of lipid profiles and Hemoglobin A1c was not conducted in the patients undergoing aortic valve replacement, because not more than three among the eight patients had these data available.

**Extended Data Table 4.** Digestion mixes for human CV tissues Digestion mixes for human aortic plaque.

| **Digestion mix I**<br>**1 mL for 200 mg tissue**<br>**Incubate at 37 °C, 300 rpm for 70 min** |  |  |
| --- | --- | --- |
| **Reagents** | **Amount** | **Supplier** |
| PBS with Ca/Mg<sup>2+</sup> | 905.3 mL | ThermoFisher |
| Collagenase I (675 U/mL) | 67.5 mL | Sigma-Aldrich |
| Collagenase XI (187,5 U/mL) | 15 mL | Sigma-Aldrich |
| Hyaluronidase (90 U/mL) | 7.5 mL | Sigma-Aldrich |
| Dnase I (90 U/mL) | 4.7 mL | Sigma-Aldrich |

| **Digestion mix II**<br>**1 mL for 200 mg tissue**<br>**Incubate at 37 °C, 300 rpm for 8 min** |  |  |
| --- | --- | --- |
| **Reagents** | **Amount** | **Supplier** |
| PBS with Ca/Mg<sup>2+</sup> | 952.5 mL | ThermoFisher |
| Collagenase II (400 U/mL) | 40 mL | Sigma-Aldrich |
| Hyaluronidase (90 U/mL) | 7.5 mL | Sigma-Aldrich |

**Digestion mixes for human heart tissue**
| **Digestion mix (1 mL for 200 mg tissue)** |  |  |
| --- | --- | --- |
| **Reagents** | **Amount** | **Supplier** |
| DMEM | 953.3 mL | ThermoFisher |
| Collagenase I (675 U/mL) | 38.7 mL | Sigma-Aldrich |
| Hyaluronidase (90 U/mL) | 3 mL | Sigma-Aldrich |
| Dnase I (90 U/mL) | 5 mL | Sigma-Aldrich |

**Extended Data Table 5.** FACS panels for human CV tissues and blood.

| **FACS panel for human aortic plaque** |  |  |  |
| --- | --- | --- | --- |
| **Target** | **Fluorochrome** | **Manufacturer** | **Dilution** |
| Viability dye | APC-Cy7 | ThermoFisher | 1:1000 |
| CD45 | PacBlue | BioLegend | 1:300 |
| CD3 | PE-ef610 | ThermoFisher | 1:300 |
| CD19 | BV605 | BioLegend | 1:300 |
| CD14 | BV711 | BioLegend | 1:300 |
| CD64 | FITC | BioLegend | 1:300 |
| MERTK | PE | BioLegend | 1:300 |

| **FACS panel for human blood (patients undergoing carotid endarterectomy)** |  |  |  |
| --- | --- | --- | --- |
| **Target** | **Fluorochrome** | **Manufacturer** | **Dilution** |
| CD45 | PacBlue | BioLegend | 1:300 |
| CD3 | PE-ef610 | BioLegend | 1:300 |
| CD4 | PerCP-Cy5.5 | BioLegend | 1:300 |
| CD8 | PECy5.5 | BioLegend | 1:300 |
| CD19 | BV605 | BioLegend | 1:300 |
| CD14 | BV711 | BioLegend | 1:300 |

| **FACS panel for human heart** |  |  |  |
| --- | --- | --- | --- |
| **Target** | **Fluorochrome** | **Manufacturer** | **Dilution** |
| Viability dye | APC-Cy7 | ThermoFisher | 1:1000 |
| CD45 | PacBlue | BioLegend | 1:300 |
| CD14 | PE | BioLegend | 1:300 |
| CD64 | FITC | BioLegend | 1:300 |
| CCR2 | APC | BioLegend | 1:300 |
| HLADR | PE-Cy7 | BioLegend | 1:300 |

| **FACS panel for human blood (patients undergoing heart surgery)** |  |  |  |
| --- | --- | --- | --- |
| **Target** | **Fluorochrome** | **Manufacturer** | **Dilution** |
| CD45 | PacBlue | BioLegend | 1:300 |
| CD3 | APC | BioLegend | 1:300 |
| CD11b | APC-Cy7 | BioLegend | 1:300 |
| HLA-DR | PE-Cy7 | BioLegend | 1:300 |
| CD15 | PerCP-Cy5.5 | BioLegend | 1:300 |
| CD16 | FITC | BioLegend | 1:300 |
| CD14 | PE | BioLegend | 1:300 |

**Extended Data Table 6.** GO pathways comprising genes for the calculation of module scores.

| Module scores | GO pathways | Number of genes |
| --- | --- | --- |
| Inflammatory response | GO:0050729 positive regulation of inflammatory response | 167 |
| Phagocytosis | GO:0050766 positive regulation of phagocytosis | 94 |
| Stimuli recognition | GO:0062208 positive regulation of pattern recognition receptor signaling pathway | 69 |
| Chemotaxis | GO:0002690 positive regulation of leukocyte chemotaxis | 112 |
| Antigen presentation | GO:0019882 antigen processing and presentation | 135 |
| Positive cell cycle regulation | GO:0045787 positive regulation of cell cycle | 367 |
| Canonical glycolysis | GO:0061621 canonical glycolysis | 17 |
| Fatty acid metabolism | GO:0045923 positive regulation of fatty acid metabolic process | 50 |
| Glutamine metabolism | GO:0006541 glutamine metabolic process | 22 |

## Reference

1. Steensma DP. Clinical consequences of clonal hematopoiesis of indeterminate potential. Blood Adv. 2018;2(22). doi:10.1182/bloodadvances.2018020222

2. Steensma DP, Bejar R, Jaiswal S, et al. Clonal hematopoiesis of indeterminate potential and its distinction from myelodysplastic syndromes. Blood. 2015;126(1):9–16. doi:10.1182/blood-2015-03-631747

3. Bick AG, Pirruccello JP, Griffin GK, et al. Genetic Interleukin 6 Signaling Deficiency Attenuates Cardiovascular Risk in Clonal Hematopoiesis. Circulation. Published online 2020:124–131. doi:10.1161/CIRCULATIONAHA.119.044362

4. Fuster JJ, MacLauchlan S, Zuriaga MA, et al. Clonal hematopoiesis associated with TET2 deficiency accelerates atherosclerosis development in mice. Science (80-). 2017;355(6327):842–847. doi:10.1126/science.aag1381

5. Sano S, Oshima K, Wang Y, et al. Tet2-Mediated Clonal Hematopoiesis Accelerates Heart Failure Through a Mechanism Involving the IL-1β/NLRP3 Inflammasome. J Am Coll Cardiol. 2018;71(8):875–886. doi:10.1016/j.jacc.2017.12.037

6. Abplanalp WT, Cremer S, John D, et al. Clonal Hematopoiesis-Driver DNMT3A Mutations Alter Immune Cells in Heart Failure. Circ Res. 2021;128(2):216–228. doi:10.1161/CIRCRESAHA.120.317104

7. Bick AG, Weinstock JS, Nandakumar SK, et al. Inherited causes of clonal haematopoiesis in 97,691 whole genomes. Nature. 2020;586(7831):763–768. doi:10.1038/s41586-020-2819-2

8. Jaiswal S, Natarajan P, Silver AJ, et al. Clonal Hematopoiesis and Risk of Atherosclerotic Cardiovascular Disease. N Engl J Med. 2017;377(2):111–121. doi:10.1056/NEJMoa1701719

9. Jaiswal S, Fontanillas P, Flannick J, et al. Age-Related Clonal Hematopoiesis Associated with Adverse Outcomes. N Engl J Med. 2014;371(26):2488–2498. doi:10.1056/nejmoa1408617

10. Svensson EC, Madar A, Campbell CD, et al. TET2 -Driven Clonal Hematopoiesis and Response to Canakinumab: An Exploratory Analysis of the CANTOS Randomized Clinical Trial. JAMA Cardiol. 2022;7(5):521–528. doi:10.1001/jamacardio.2022.0386

11. Genovese G, Kähler AK, Handsaker RE, et al. Clonal Hematopoiesis and Blood-Cancer Risk Inferred from Blood DNA Sequence. N Engl J Med. 2014;371(26):2477–2487. doi:10.1056/nejmoa1409405

12. Tall AR, Fuster JJ. Clonal hematopoiesis in cardiovascular disease and therapeutic implications. Nat Cardiovasc Res. 2022;1(2). doi:10.1038/s44161-021-00015-3

13. Liu W, Yalcinkaya M, Maestre IF, et al. Blockade of IL-6 signaling alleviates atherosclerosis in Tet2-deficient clonal hematopoiesis. Nat Cardiovasc Res. 2023;2(6):572–586. doi:10.1038/s44161-023-00281-3

14. Dorsheimer L, Assmus B, Rasper T, et al. Association of Mutations Contributing to Clonal Hematopoiesis with Prognosis in Chronic Ischemic Heart Failure. JAMA Cardiol. 2019;4(1):25–33. doi:10.1001/jamacardio.2018.3965

15. Arends CM, Liman TG, Strzelecka PM, et al. Associations of clonal hematopoiesis with recurrent vascular events and death in patients with incident ischemic stroke. Blood. 2023;141(7):787–799. doi:10.1182/blood.2022017661

16. Kittang AO, Kordasti S, Sand KE, et al. Expansion of myeloid derived suppressor cells correlates with number of T regulatory cells and disease progression in myelodysplastic syndrome. Oncoimmunology. 2016;5(2). doi:10.1080/2162402X.2015.1062208

17. Arends CM, Galan-Sousa J, Hoyer K, et al. Hematopoietic lineage distribution and evolutionary dynamics of clonal hematopoiesis. Leukemia. 2018;32(9):1908–1919. doi:10.1038/s41375-018-0047-7

18. Bajpai G, Schneider C, Wong N, et al. The human heart contains distinct macrophage subsets with divergent origins and functions. Nat Med. 2018;24(8):1234–1245. doi:10.1038/s41591-018-0059-x

19. Mas-Peiro S, Hoffmann J, Fichtlscherer S, et al. Clonal haematopoiesis in patients with degenerative aortic valve stenosis undergoing transcatheter aortic valve implantation. Eur Heart J. 2020;41(8):933–939. doi:10.1093/eurheartj/ehz591

20. Dederichs T-S, Ehlert C, Becker H, Pfeifer D, Bode C, Hilgendorf I. Chip mutations mediate human atherosclerosis by activating monocyte pro-inflammatory pathways without evidently promoting monocyte chemotaxis. Atherosclerosis. 2021;331:e12. doi:10.1016/j.atherosclerosis.2021.06.041

21. Swirski FK, Nahrendorf M. Leukocyte behavior in atherosclerosis, myocardial infarction, and heart failure. Science (80-). 2013;339(6116). doi:10.1126/science.1230719

22. Robbins CS, Hilgendorf I, Weber GF, et al. Local proliferation dominates lesional macrophage accumulation in atherosclerosis. Nat Med. 2013;19(9). doi:10.1038/nm.3258

23. Härdtner C, Kornemann J, Krebs K, et al. Inhibition of macrophage proliferation dominates plaque regression in response to cholesterol lowering. Basic Res Cardiol. 2020;115(6):1–19. doi:10.1007/s00395-020-00838-4

24. Nam AS, Dusaj N, Izzo F, et al. Single-cell multi-omics of human clonal hematopoiesis reveals that DNMT3A R882 mutations perturb early progenitor states through selective hypomethylation. Nat Genet. Published online 2022. doi:10.1038/s41588-022-01179-9

25. Bouzid H, Belk JA, Jan M, et al. Clonal hematopoiesis is associated with protection from Alzheimer’s disease. Nat Med. 2023;29(July). doi:10.1038/s41591-023-02397-2

26. Sager RA, Khan F, Toneatto L, et al. Targeting extracellular Hsp90: A unique frontier against cancer. Front Mol Biosci. 2022;9. doi:10.3389/fmolb.2022.982593

27. Choudhury A, Bullock D, Lim A, et al. Inhibition of HSP90 and Activation of HSF1 Diminish Macrophage NLRP3 Inflammasome Activity in Alcohol-Associated Liver Injury. Alcohol Clin Exp Res. 2020;44(6). doi:10.1111/acer.14338

28. Leclerc E, Fritz G, Weibel M, Heizmann CW, Galichet A. S100B and S100A6 differentially modulate cell survival by interacting with distinct RAGE (receptor for advanced glycation end products) immunoglobulin domains. J Biol Chem. 2007;282(43). doi:10.1074/jbc.M703951200

29. Rocha VZ, Libby P. Obesity, inflammation, and atherosclerosis. Nat Rev Cardiol. 2009;6(6). doi:10.1038/nrcardio.2009.55

30. Libby P, Buring JE, Badimon L, et al. Atherosclerosis. Nat Rev Dis Prim. 2019;5(1):56. doi:10.1038/s41572-019-0106-z

31. Tamargo IA, Baek KI, Kim Y, Park C, Jo H. Flow-induced reprogramming of endothelial cells in atherosclerosis. Nat Rev Cardiol. Published online 2023. doi:10.1038/s41569-023-00883-1

32. Simón Chica A, M. Wülfers E, Kohl P. Non-myocytes as electrophysiological contributors to cardiac excitation and conduction. Am J Physiol Circ Physiol. 0(0):null. doi:10.1152/ajpheart.00184.2023

33. Lother A, Kohl P. The heterocellular heart: identities, interactions, and implications for cardiology. Basic Res Cardiol. 2023;118(1):30. doi:10.1007/s00395-023-01000-6

34. Peet C, Ivetic A, Bromage DI, Shah AM. Cardiac monocytes and macrophages after myocardial infarction. Cardiovasc Res. 2020;116(6). doi:10.1093/CVR/CVZ336

